# A chemical structure and machine learning approach to assess the potential bioactivity of endogenous metabolites and their association with early-childhood hs-CRP levels

**DOI:** 10.1101/2023.11.15.567095

**Authors:** Mario Lovrić, Tingting Wang, Mads Rønnow Staffe, Iva Šunić, Kristina Časni, Jessica Lasky-Su, Bo Chawes, Morten Arendt Rasmussen

## Abstract

Metabolomics has gained much attraction due to its potential to reveal molecular disease mechanisms and present viable biomarkers. In this work we used a panel of untargeted serum metabolomes in 602 childhood patients of the COPSAC2010 mother-child cohort. The annotated part of the metabolome consists of 493 chemical compounds curated using automated procedures. Using predicted quantitative-structure-bioactivity relationships for the Tox21 database on nuclear receptors and stress response in cell lines, we created a filtering method for the vast number of quantified metabolites. The metabolites measured in children’s serums used here have predicted potential against the chosen target modelled targets. The targets from Tox21 have been used with quantitative structure-activity relationships (QSARs) and were trained for ∼7000 structures, saved as models, and then applied to 493 metabolites to predict their potential bioactivities. The models were selected based on strict accuracy criteria surpassing random effects. After application, 52 metabolites showed potential bioactivity based on structural similarity with known active compounds from the Tox21 set. The filtered compounds were subsequently used and weighted by their bioactive potential to show an association with early childhood hs-CRP levels at six months in a linear model supporting a physiological adverse effect on systemic low-grade inflammation. The significant metabolites were reported.

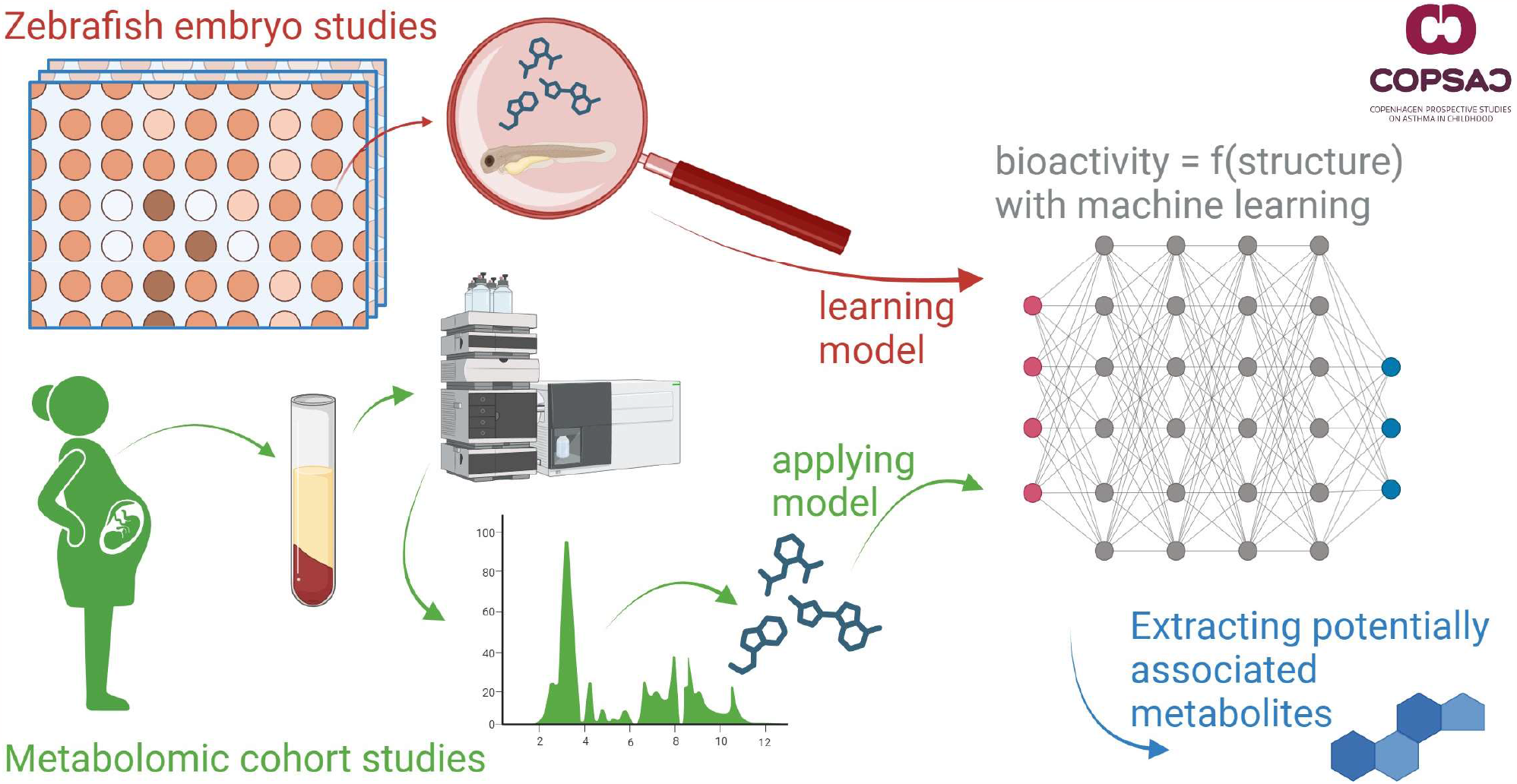

## 1 Introduction

Metabolomics aims to capture a broad range of small molecules, either quantitatively by measuring their concentration or qualitatively by identifying their presence and structure, which are essential intermediates and end products of metabolism in organisms. One can analyze pathological processes underlying a disease or physiological state of interest using rich metabolomics data, e.g., obtaining biomarkers (1) or predicting responses to therapy (2,3). Furthermore, one can use the metabolome for phenotyping organisms or diseases (4). Metabolomic instruments provide rapid, cost-effective, and sensitive results. Yet, with appropriate analysis, the data they produce is meaningful. It’s essential to interpret this data to build biochemical pathways and comprehend their interactions in healthy and diseased conditions (5). Liquid chromatography-mass - mass spectrometry (LC-MS) as one of the most commonly used approaches for metabolomics study can achieve a high number of annotations (6), which is also the technique used in this study. A metabolome is a single snapshot of the organism’s pathophysiological state (7) and must be treated cautiously when interpreting and trying to understand underlying causality. Another significant predictor of health risks is the assessment of levels of C-reactive protein with high-sensitivity methods (hs-CRP), considered generally a marker of systemic low-grade inflammation. Elevated hs-CRP has been shown in early-onset diseases and has been associated with an increased prevalence of overweight and obesity in children (8,9). Studies have shown elevated hs-CRP is a biomarker of childhood asthma(10)and allergy (11) but also associated with ADHD development (12). Hence, hs-CRP is a valuable tool for understanding childhood obesity risk, asthma, allergy, and overall health status, and elevated hs-CRP is interpreted as a physiological state of inflammation in this study. Inflammation is assumed to be related to metabolic processes and should hence be reflected in metabolome by, e.g., other biomarkers like the previously presented GlycA (13). The leading assumption in this work is that one can use chemical information of endogenous or exogeneous metabolites to relate them to physiological states. This can be processed using quantitative structure-activity relationships (QSARs). QSARs are based on mapping a feature space (X), calculated from chemical structures, on a target chemical or biological activity (y)(14,15). In reverse, after developing a QSAR, one can predict biological activity for a specified target for an (in the model) unseen chemical structure, given some chemical similarity to previously utilized chemical space. To achieve this with metabolites, one must know their structures. Biological activities of interest herein are activation or inhibition of specific biochemical pathways. Hence, to understand the models created and differentiate the concept from an environmental perspective, compounds analyzed against targets will be deemed “bioactive” or even “toxic.” The selected dataset for modeling stems from the Tox21 compound library (16,17) and includes 12 bioactivity endpoints, five related to stress response (SR) and seven to nuclear receptor (NR) panels. NRs are a class of transcription factors involved in regulating gene expression. When a chemical compound binds to an NR, it induces a conformational change that allows the NR to interact with coactivators or corepressors, which regulate gene transcription, finally leading to detrimental effects on the organism. The SR pathways are activated in response to various environmental and cellular stresses, including oxidative stress, DNA damage, and protein misfolding. This work aims to filter metabolites from many compounds to inspect associations with physiological conditions. Filtering is based on structural similarities of metabolites to known bioactive compounds, which can be potentially bioactive in unspecified pathways. Hence, if dysregulated, endogenous compounds could be associated with physiological states. Even though there is not much research on the endobiotic effects and associations, there are indications for such. Elevated cortisol concentrations has been associated to psychiatric diseases (18,19). Other metabolites like phenylalanine are known toxicants in conditions such as phenylketonuria (20). Research also suggests toxic intermediates in metabolomic pathways (Lee et al., 2020) can contain reactive functional groups and be leveraged for, e.g., cancer therapy. A set of QSAR models is developed in this work against the Tox21 biological target and used to predict potential outcomes for each of the quantified metabolites. The predicted outcomes are used then as filters to select potentially bioactive compounds that could trigger the given Tox21 pathways. The filtered metabolites were evaluated based on their abundance and bioactivity against levels of hs-CRP in the children at six months of age.

## 2 Materials and methods

### 2.1 Cohort metabolomics and clinical outcomes

#### 2.1.1 Cohort

The COPSAC2010 cohort comprises a non-selected group of mother-child pairs, including 738 pregnant women and their 700 children. The women were recruited between gestational weeks 22-26 during their first pregnancy examinations. Of the 738 pregnant women (with an average age of 32.3±4.3 years at the time of their child’s birth), 700 children were enrolled in the study. The children visited the clinic for regular visits since then. Gestational age was determined using routine pregnancy care ultrasonography. The COPSAC2010 study participants included infants delivered pre-term and post-term (30-42 weeks). During the third trimester, the women took part in a double-blind, randomized controlled trial with a factorial design, receiving either high-dose vitamin D (2800 IU/day) or standard dose (400 IU/day) (21), and either 2.4 g n-3 long-chain polyunsaturated fatty acid (LCPUFA, 55% (w/w) 20:5(n-3) eicosapentaenoic acid (EPA) and 37% (w/w) 22:6(n-3) docosahexaenoic acid (DHA)) or placebo (72% (w/w) n-9 oleic acid and 12% (w/w) n-9 linoleic acid) (22). Women with endocrine, heart, or kidney diseases or a daily vitamin D intake above 600 IU/day were excluded. Children with a gestational age of less than 32 weeks were also excluded. The trial received approval from the National Committee on Health Research Ethics (H-B-2008-093) and the Danish Data Protection Agency (2015-41-3696). Both parents provided oral and written informed consent before enrolment.

#### 2.1.2 Blood collection and metabolites quantification

Blood samples were taken from the mothers at week 24 of pregnancy, one week postpartum, and six months from children. Blood was collected in an ethylenediaminetetraacetic acid (EDTA) tube (lithium-heparin tube for child samples at 18 months) and left at room temperature for 30 minutes before being centrifuged for 10 minutes at 4000 rpm. The supernatant was collected and stored at -80 °C for future analysis (23). The sample preparation (Ultra High-Performance Liquid Chromatography—Tandem Mass Spectrometry), UHPLC-MS/MS analysis, and quality control for blood metabolomic profiling of mothers have been previously described (24). Metabolon, Inc. in Morrisville, NC, USA, conducted untargeted plasma metabolomic analysis on mothers’ and children’s blood. They used an ACQUITY UHPLC QExactive™ Hybrid Quadrupole-Orbitrap™ mass spectrometer interfaced with a heated electrospray ionization source operated at 35,000 mass resolution. The processed samples were analyzed on four platforms, including UHPLC-ESI(+)MS/MS optimized for hydrophilic compounds, UHPLC-ESI(+)MS/MS optimized for hydrophobic compounds, reverse phase UHPLC-ESI(−)MS/MS using basic optimized conditions, and HILIC/UHPLC-ESI(−)MS/MS. Metabolites were identified based on three matching criteria: retention time/index range, mass accuracy (±10 ppm), and MS/MS spectra.

#### 2.1.3 Assessment of hs-CRP levels

Children at six months old had blood drawn from a cubital vein into an EDTA tube. The samples were centrifuged to separate plasma and cells and stored at −80 °C until analysis. After the samples were thawed, the hs-CRP levels were measured using a high-sensitivity electrochemiluminescence-based assay from MesoScale Discovery. Duplicate measurements were taken and analyzed using the Sector Image 2400 A from Meso Scale Discovery in Gaithersburg, MD. The lower limit of detection for CRP was 0.007 ng/mL (25).

### 2.2 Data preparation and model building

#### 2.2.1 Data preparation for metabolites

The metabolites with more than 33% missing values were removed from the analysis. Those with missing values were imputed with 1/10 of the minimum contraction per metabolite (under the assumption to correspond to the detection limit). The metabolites were then scaled to a 0-1 (min-max scaling). A further cleaning step was the removal of low-variance metabolites, i.e., the lowest 10% of metabolites by variance. The last step was removing highly correlated metabolites with above 90% Pearson correlation. In total, 517 compounds or 602 mom-child pairs, i.e., metabolites names, were converted to SMILES encodings of molecules, resulting in a final data set for the analysis.

#### 2.2.2 Bioactivity assessment pipeline

The model training and application pipeline is presented in Figure 1. The pipeline starts with data extraction and preparation for building QSAR models (blue and yellow rectangles in the figures), described in Sections 2.2.2. and 2.2.3. Once data is prepared, models are built using Random Forests and hyperparameter optimization (Section 2.2.4.) on features generated from chemical structures. These models can predict bioactivity against biological targets such as the 12 given in Section 2.2.2. Once the models are generated and validated, the final step is to predict bioactivities for the 12 targets per quantified metabolite and patient. The predictions then enter further statistical analysis to reveal patterns in clinical outcomes.

**Figure 1.**
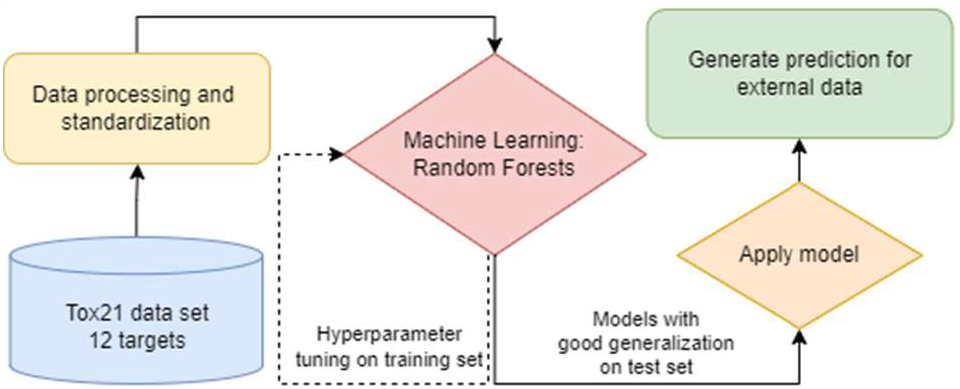
The modeling pipeline in this study.

#### 2.2.3 Bioactivity data

The chosen bioactivities for this task stem from the Tox21 compound library (16,17) provided by the U.S. Environmental Protection Agency (EPA). In the Tox21 program, these include the androgen receptor (AR), estrogen receptor (ER), progesterone receptor (PR), aromatase receptor, peroxisome proliferator-activated receptor gamma (PPAR), and the aryl hydrocarbon receptor (AhR). The compounds in the library were tested against their ability to interact with various SR pathways, including antioxidant-responsive elements (ARE), p53 tumor proteins, mitochondrial membrane potential (MMP), proteins involved in DNA damage repair (ATAD5), heat shock factor response elements (HSE). Numerous studies have been conducted on this dataset, and the outcomes have been reported in various reports (26–29). Hence, it represents a baseline data set for building bioactivity QSARs since it is imbalanced, chemically diverse, and one of the more significant publicly available (∼10k compounds). Because of the challenges presented by this dataset, it has been extensively studied in cheminformatics research. In machine learning methods (26,29), class balancing methods (28,30), and testing of different chemical representations (27) and multitask/deep learning (29). The raw data was pre-processed as it contains duplicated structures considering the active or organic part of the molecules. This was also reported earlier (28). Molecules with invalid structural identifiers were removed, and those that were valid were converted to their canonical SMILES (31). Duplicates were either removed by IDs or SMILES. The inorganic compounds, metal-containing compounds, and fragments were also removed, inspired by procedures (28,32) to keep the active parts of the molecules. Once structures were ready, molecular descriptors Morgan fingerprints (FPR) (33) and 200 molecular descriptors were calculated for the 8,314 structures using the RDKit library (34). To reduce possible bit collision in fingerprints (35,36), the fingerprint vectors were set to 5,120 bits and a radius of 2. The Python scripts used for this work were published with (37).

#### 2.2.4 Machine learning model

Machine learning models were trained using Python (www.python.org) and its library sci-kit-learn (38) based on previous works (37,39). Due to class imbalance in the Tox21 set, model penalization and optimization techniques were used to improve classification quality. Before model training, the data with the 12 bioactivity endpoints was split into two random subsets of 80% and 20% per endpoint individually. The penalty in scoring during hyperparameter optimization was based on the Matthews correlation coefficient (MCC) (40,41) defined by equation 1, where TP (True Positive), TN (True Negative), FN (False Negative), FP (False Positive) are the elements of the confusion matrix.^1^

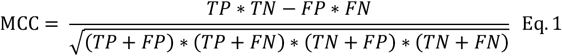

Bayesian hyperparameter optimization (BO) was used for hyperparameter optimization (41,42) through a 10-fold cross-validation and penalizing the procedure by MCC iterating it 20 times and returning a set of “optimal” parameters. One of the essential model parameters was “class weight,” which contributed to classifying this imbalanced data set. During model training on the train set, feature selection using permutation importance was applied in the following steps in previous work (41).

### 2.3 Association testing and toxic unit approach

To evaluate the association to physiological conditions the toxic unit (TU) approach was utilized (43). TU is defined as the ratio of chemical compound concentration (ci) and the selected exposure-based toxicity value (e.g., LC50). Herein, a new approach is derived, namely the bioactive potential (*BiP*) in Eq.1. *BiP* is hence a product of the metabolite’s concentration (*i*) (relative peak area) and the probability of it being active (0-1) against a target (*t*). The higher the concentration and the likelihood of being bioactive, the more potent we assume the compound to be against the given targets.

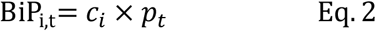

For each child (*c*) in the cohort, *BiP* was calculated per metabolite and target and then summed and log-transformed *sBIP* using Eq.2, where *T* is the number of targets, yielding in one value per child.

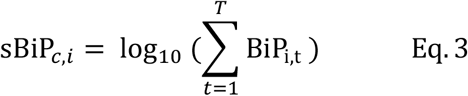

The associations were inspected by means of linear regression.

## 3 Results and discussion

### 3.1 QSAR model results

Eight out of twelve models yielded results above random classification (MCC CV and MCC Test > 0.3), namely for the following targets: SR-ATAD5, NR-AR-LBD, NR-AR, NR-ER-LBD, NR-ER, SR-ARE, NR-AhR, SR-MMP. These models were then selected for further processing. While the algorithm trained for both fingerprints and descriptor set separately, each time, the descriptors yielded better results in this setting over fingerprints.

### 3.2 Predicting metabolites target activity

The selected eight models were saved to persistent storage and applied to yield bioactivity for the 493 annotated metabolites. This resulted in a table of 493 metabolites x eight columns, namely SR-ATAD5, NR-AR-LBD, NR-AR, NR-ER-LBD, NR-ER, SR-ARE, NR-AhR, and SR-MMP. A snippet of the results is given in Table 2 to ease understanding of the model results. The results in the table are presented as probabilities instead of a binary result (bioactive or not), where the convention is that a probability above 0.5 (p > 0.5) is considered bioactive. One can observe from the table that the probabilities show a broad range across targets. Those marked with an asterisk would be deemed bioactive towards the target since their values are above 0.5.

**Table 1.** Model results for the 12 targets from the Tox21 data set.

| Endpoint | MCC Train | Balanced Accuracy Train | MCC CV | MCC Test | Balanced Accuracy Test |
| --- | --- | --- | --- | --- | --- |
| SR-HSE | 0.47 | 0.82 | 0.27 | 0.23 | 0.69 |
| *NR-AR | 0.63 | 0.95 | 0.58 | 0.7 | 0.97 |
| *SR-ARE | 0.77 | 0.94 | 0.35 | 0.34 | 0.76 |
| NR-Aromatase | 0.6 | 0.88 | 0.2 | 0.15 | 0.63 |
| *NR-ER-LBD | 0.62 | 0.89 | 0.47 | 0.56 | 0.84 |
| *NR-AhR | 0.73 | 0.9 | 0.51 | 0.45 | 0.78 |
| *SR-MMP | 0.83 | 0.91 | 0.59 | 0.58 | 0.79 |
| *NR-ER | 0.64 | 0.91 | 0.41 | 0.41 | 0.83 |
| NR-PPAR-gamma | 0.59 | 0.92 | 0.17 | 0.09 | 0.6 |
| SR-p53 | 0.41 | 0.63 | 0.25 | 0.24 | 0.58 |
| *SR-ATAD5 | 0.52 | 0.76 | 0.3 | 0.31 | 0.66 |
| *NR-AR-LBD | 0.61 | 0.81 | 0.58 | 0.55 | 0.82 |

**Table 2.** A snippet of probabilities of metabolites being active towards the eight models (columns)

|  | SR-ATAD5 | NR-AR-LBD | NR-AR | NR-ER-LBD | NR-ER | SR-ARE | NR-AhR | SR-MMP |
| --- | --- | --- | --- | --- | --- | --- | --- | --- |
| cortisol | 0.12 | *0.97 | *0.95 | 0.07 | 0.35 | 0.2 | 0 | 0.13 |
| cortisone | 0.13 | *0.96 | *0.82 | 0.04 | 0.36 | 0.24 | 0.02 | 0.19 |
| androsterone sulfate | 0.12 | *0.88 | *0.67 | 0.43 | *0.59 | 0.48 | 0.02 | 0.33 |
| dehydroepiandrosterone sulfate | 0.13 | *0.92 | *0.66 | 0.46 | *0.67 | *0.54 | 0.01 | 0.29 |
| 16a-hydroxy DHEA 3-sulfate | 0.14 | *0.93 | *0.63 | 0.37 | *0.57 | 0.42 | 0.02 | 0.44 |

A mask was applied to filter out all “nonactive” results with values below 0.5 in the full results. In total, 51 metabolites showed at least one probability in one model being active above 0.5. A complete list of active metabolites is given in Table 3.

**Table 3.**
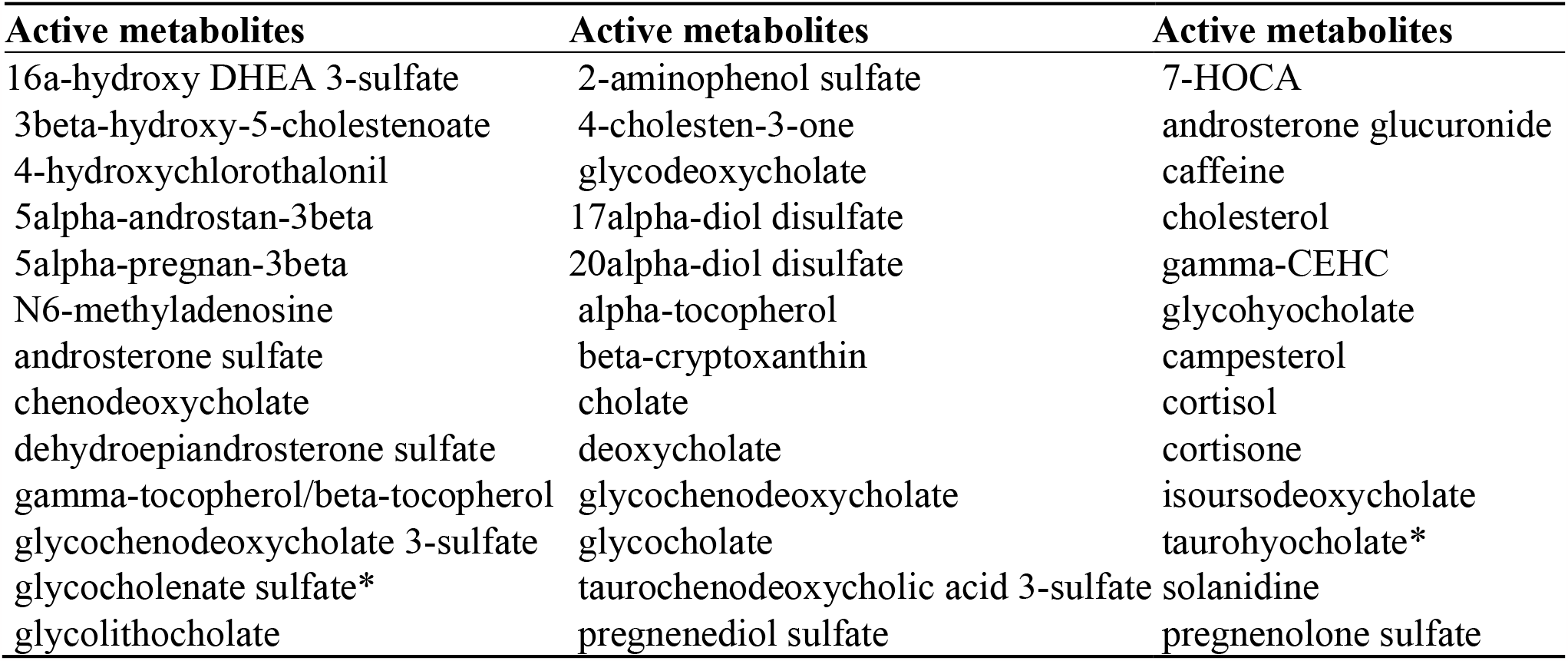
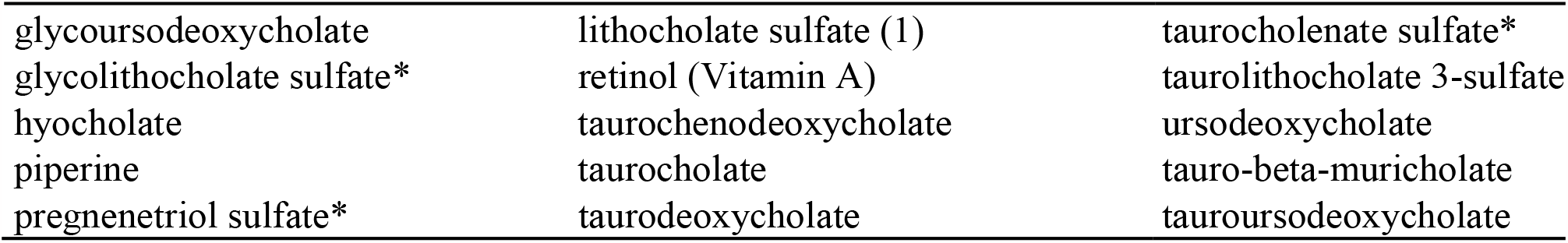
A list of active metabolites given the pre-trained models.

### 3.3 Association with hs-CRP

For each child (*c*) in the cohort, *BiP* was calculated per metabolite and target and then summed and log-transformed *sBIP* using Eq.2. Once a table was obtained with *sBiP* per child and metabolite, the potentially bioactive part of the metabolome was associated with levels of hs-CRP. The table was scaled priorly, and metabolites were decorrelated, reducing their number from 51 to 40. The hs-CRP levels were logarithmed, and missing values were imputed by median values. The associations were then evaluated in an linear regression. Metabolites amongst the 40 metabolites with an association with hs-CRP filtered for an regression coefficient (estimate) with a p-value below 0.001 are shown in Table 4. The results show negative significant coefficients for cortisone, retinol, and 3beta-hydroxy-5-cholestenoate, while the cortisol and glycocholenate sulfate resulted in positive coefficients.

**Table 4.** Results of the association testing against hs-CRP.

| metabolite | Super_pathway | Sub_pathway | coefficient | p* |
| --- | --- | --- | --- | --- |
| cortisone | Lipid | Corticosteroids | -0.91 | 0.000 |
| retinol (Vitamin A) | Cofactors and Vitamins | Vitamin A Metabolism | -1.39 | 0.000 |
| cortisol | Lipid | Corticosteroids | 1.026 | 0.000 |
| 3beta-hydroxy-5-cholestenoate | Lipid | Sterol (or primary bile acid biosynthesis) | -0.64 | 0.000 |
| glycocholenate sulfate* | Lipid | Secondary Bile Acid Metabolism | 0.581 | 0.002 |
| intercept |  |  | 2.976 | 0.000 |

## 4 Discussion

### 4.1 Discussion of results

hs-CRP is a well-known biomarker in the field of medicine and clinical diagnostics, serving as a powerful tool for assessing low-grade or chronic inflammation (44,45). We have previously shown that systemic low-grade inflammation assessed by hs-CRP levels in pregnant mothers and their children are correlated independent of anthropometrics, environmental exposures(25). Elevated levels of hs-CRP are associated with an increased risk of cardiovascular disease (46), inflammatory bowel disease (IBD)(47), depression (48), ADHD (12) decreased lung function in childhood (49), allergic sensitization (11), early-life airway microbiota (50), and childhood asthma (51). Metabolites serve as the ultimate effectors of cellular processes and represent the penultimate step in the progression to the phenotype. This underscores the importance of investigating the association between CRP and metabolites to gain a deeper understanding of how CRP is reflected in the metabolome. In our research, we developed a series of QSAR models targeting the Tox21 cellular endpoints. These models were employed to forecast potential outcomes against such targets for each of the quantified metabolites within the COPSAC2010 cohort. The selection of these specific metabolites was based on their abundance and bioactivity in relation to high-sensitivity C-reactive protein (hs-CRP) levels in children at six months of age. First and foremost, the model we developed has achieved reasonable generalization, as indicated by the prediction results. The balanced accuracy of our models for the selected eight endpoints consistently exceeded 0.6 in both the training and test sets, and the Matthews Correlation Coefficients (MCC) consistently exceeded 0.3. These metrics demonstrate the robustness of our models for making accurate predictions in the following sections. Using the developed models, we identified 40 out of the 493 metabolites in our cohort that showed bioactivity in at least one of the eight models. We test the BiP of these 40 metabolites in a linear model against the levels of high-sensitivity hs-CRP in children at six months of age. Notably, we identified five metabolites of particular interest, each demonstrating regression coefficients (estimates) with a p-value below 0.001. Among these five metabolites, cortisone and cortisol are both belong to corticosteroid hormones, and cortisone is released by the adrenal gland in response to stress (51). We observed positive association between cortisol and hs-CRP and negative association between CRP and cortisone. One of cortisone’s effects on the body, is the suppression of the immune system due to a decrease in the function of the lymphatic system (53), which is consistent with our findings. It was proven in Shimba’s study (54) that corticosteroids are strong inhibitors of inflammatory corneal lymphangiogenesis, with significant differences between various corticosteroids in terms of their antilymphangiogenic potency. The main mechanism seems to be through the suppression of macrophage infiltration, proinflammatory cytokine expression, and inhibition of proliferation of lymphatic endothelial cells. In study of Rueggeberg et al., it was observed that 2-year increases in diurnal cortisol secretion were significantly associated with higher levels of CRP at 6-year follow-up (55).These findings may difficult to reconcile because cortisol generally has anti-inflammatory properties. However, sustained exposure to high levels of cortisol may render innate immune cells partially resistant to glucocorticoid inhibition, allowing inflammation to escape normal regulatory controls (56,57). This could be an explanation for the apparent positive correlation between high CRP and cortisol. The coefficients of cortisol and cortisone show an opposite direction (1.026 vs -0.91) with both being significant. The balance between serum cortisol and cortisone is a known phenomenon and hence often a ratio is used (cortisol/cortisone) with cortisol being the more active one. Examples of altered rations comparing to normal can occur during acute-phase responses (58), in obesity related issues (59), and tuberculosis (60). The relationship in this work may subject to various factors, including the presence of chronic stress, individual differences, and the balance between pro-inflammatory and anti-inflammatory processes in the body. A positive association between cortisol and CRP and a negative association between cortisone and CRP can be explained by the complex and dynamic nature of the body’s stress and inflammatory responses. There are situations where cortisol can have pro-inflammatory effects under chronic stress (61). Chronic stress can lead to dysregulation of the hypothalamic-pituitary-adrenal axis, and subsequently results in prolonged elevated cortisol levels (62) These prolonged high levels of cortisol can induce inflammation and stimulate the production of inflammatory mediators, like CRP. In this scenario, a positive association between cortisol and CRP can be observed. Retinol (Vitamin A) exhibited the most robust negative association with hs-CRP among these five notable metabolites in our predictive analysis. In the study by Dios et al. (63), it was investigated that 6-8-year old children in the highest hs-CRP group (hs-CRP ≥ 0.60 mg/dL) displayed significantly lower levels of retinol compared to those in the lower hs-CRP groups. It is hypothesized that retinol binding to retinol-binding protein (RBP) plays a role in the association between retinol and CRP in children. From a biological perspective, retinol is transported in the blood while bound to RBP, and its concentration is tightly regulated by RBP (64). Additionally, a correlation between RBP4 and CRP levels has been described in school-aged obese children(65). Our findings align with previous research, and we have further confirmed the strong correlation between retinol and CRP in a larger, controlled cohort at 6 months child. 3beta-hydroxy-5-cholestenoate (3-BH5C) belongs to the class of monohydroxy bile acids, alcohols, and derivatives. This compound is a component of the primary bile acid biosynthesis pathway and has been recognized as a metabolite capable of predicting gut microbiome Shannon diversity (66). Shannon diversity serves as a potential marker for overall microbiome health. In our results, we identified a strong negative association between 3-BH5C and CRP. To the best of our knowledge, this is the first time a negative correlation between 3-BH5C and CRP has been observed, suggesting that the microbiome health of children, as indicated by the marker 3-BH5C, may be linked to inflammation. A prior study conducted in the Northern Finland Birth Cohort 1966 (NFBC1966) and the TwinsUK cohort demonstrated that higher CRP levels were associated with lower alpha diversity of gut microbiome measures (67). This provides further support for the credibility of our findings. Another bile acid metabolite, glycocholenate sulfate, classified as a conjugated bile acid (CBA), has been identified as a significant biomarker for CVD risk in a cohort that included 1919 African-American participants in the Atherosclerosis Risk study (68). CBA profiles are increasingly recognized as crucial signaling molecules intricately involved in mammalian cholesterol and lipid metabolism, glucose homeostasis, thermogenesis, inflammation, and intestinal function (69). In our study, it was further confirmed that CBA, represented by glycocholenate sulfate, is strongly associated with CRP levels in children at 6 months.

### 4.2 Limitations

There are several limitations to consider when interpreting the results. First, the metabolite concentrations were measured semi-quantitatively, which may introduce some degree of measurement error. Secondly, the bioactivity model-based approach used in this study may introduce error propagation and be limited by noise (70) since a) bioactivity data has a measurement error and b) the models come with limited accuracy. Furthermore, it is essential to note that the ligand-receptor interactions are complex and may not be fully captured by the model-based approach, a known limitation of QSAR (71). While efforts were made to incorporate as much information as possible into the models, the true nature of these interactions may be more nuanced than what was captured in this study. Finally, there is no consensus on how to clean data for modeling; hence, the data-cleaning process used in this study may have needed to be more rigorous and lost some information, which is further driven by not annotated metabolites. There is also the question of causality since the exact drivers of metabolic pathways still need to be completely revealed. Future work should address some of these limitations. Compounds were excluded in the data processing; however, their effects might be more complex and synergy.

## 5 Conclusion

We developed a series of QSAR models targeting the Tox21 biological endpoint and used them to predict potential outcomes for each of the quantified metabolites within the COPSAC2010 cohort. Our analysis unveiled five metabolites of particular interest due to their robust association with hs-CRP levels. We observed the significant influence of CRP on corticosteroid hormones, bile acid metabolites, and vitamin A. Association to corticosteroid hormones is supported by literature and hence shows this data-driven and chemistry-informed approach meaningful as it leads to known findings without the addition of prior medical knowledge. Our findings regarding bile acid metabolites are novel to the best of our knowledge but are supported by known findings regarding the relationship of microbiome driven gut-health and the level of CRP. Even though causality hasn’t been research in this paper, we show that there is a relationship between metabolites and CRP. This has the potential to understanding biological pathways but also in clinical settings where supplementation with e.g., vitamin A and the improvement of gut health might play a role in early childhood

## 7 Author Contributions

ML - Data curation, Methodology, Conceptualization, Software, Visualization, Writing – original draft; TW – Validation, Investigation, Writing – original draft; MRS - Data curation, Formal analysis, Software; IŠ - Data curation, Investigation, Writing – original draft; KP - Data curation, Investigation, Software, Writing – original draft; JLS - Methodology, Conceptualization, Writing – review & editing; BC - Funding acquisition, Conceptualization, Project administration, Supervision, Writing – review & editing; MAR - Funding acquisition, Conceptualization, Supervision, Writing – review & editing

## 9 Acknowledgments

We express our deepest gratitude to the children and families of the COPSAC 2010 cohort study for all their support and commitment.

## Footnotes

1 https://en.wikipedia.org/wiki/Confusion_matrix

